# Production of recombinant human acid α-glucosidase with mannosidic N-glycans in α-mannosidase I mutant rice cell suspension culture

**DOI:** 10.1101/2022.02.07.479152

**Authors:** Jae-Wan Jung

## Abstract

High-mannose glycans, containing 6-9 mannose residues, are favorable for *in vitro* mannose phosphorylation to produce Mannose-6-phosphate (M6P) residues. These are important for lysosomal enzyme targeting and essential for uptake. Golgi α-mannosidase-I mediates mannose trimming in the N-glycosylation pathway. In this study, I used mutant rice with a T-DNA insertion in the Os04g0606400 (Δ*α-man*I) gene to produce recombinant human acid α-glucosidase (rhGAA) for enzyme replacement therapy to treat Pompe disease. After characterization of Δ*α-manI* mutant rice, the gene encoding rhGAA was introduced into mutant calli by *Agrobacterium*-mediated transformation. Integration of the target sequence into the rice genome was detected by genomic DNA PCR. Expression of rhGAA from Δ*α-man*I mutant (Δ*α-man*I-GAA) and its secretion into culture media were confirmed by western blot analysis. The N-glycosylation pattern of purified Δ*α-man*I-GAA was analysed by MALDI-TOF mass spectrometry. This showed that the high-mannose type N-glycan with man8 (63.8%) was the most abundant form, followed by Man7 (22.5%), GnGnXF3 (6.7%), Man6 (5.4%), and Man5 (1.6%). The results suggest the successful production of rhGAA with N-glycan containing 6-8 mannose residue in the Δ*α-man*I mutant rice cell suspension culture.

## Introduction

Pompe disease is an inherited genetic metabolism disorder and one of several lysosomal storage diseases. Deficiency of recombinant human acid α-glucosidase (rhGAA) causes accumulation of glycogen in muscle cells and varying levels of organ damage and rates of progression to death. Enzyme replacement therapy (ERT) is the current treatment and the enzyme carrying mannose-6-phosphate (M6P) on its N-glycan is taken up into target lysosome by binding to the mannose-6-phosphate receptor [1–3].

Early attempts to use rhGAA purified from Aspergillus niger[4] or human placenta[5] for ERT were unsuccessful, and it might be because of low enzyme dosage, disease stage, and the lack of M6P residues for uptake into muscle cells[1]. Use of rhGAA from the milk of transgenic rabbits showed improved respiratory function and restoration of some muscle function in infants [2, 6]. rhGAA from Chinese hamster ovary (CHO) cells led to the development of a drug called alglucosidase alfa (Myozyme©), and approved by the US Food and Drug Administration (FDA) for the treatment of infants and children with Pompe disease[1]. Uptake efficiency of rhGAA into target muscle cell is highly dependent on the number of M6P moieties and CHO cell-derived rhGAA has 1-2 N-glycans with M6P out of 7 potential N-glycosylation site. More recently, chemical conjugation of M6P onto the periodate-treated rhGAA (neoGAA) was shown to improve its uptake [7–10]. Neo-GAA showed highly improved clearance of lysosomal glycogen in the muscle of GAA knockout mice compared with the unmodified enzyme. Nevertheless, high cost of the ERT enzymes derived from CHO cells approved for treating lysosomal storage disease[11, 12] creates a cost burden for patients (approximately 500,000 USD per year for the life of the adult patient)[13].

Plant cell culture systems have many advantages over animal or bacterial cell culture, such as cost effectiveness, safety and eukaryotic protein processing [14–16]. The simplicity of the culture media, composed of simple carbon sources and hormones, can highly reduce production cost. Although differences in the glycosylation and cellular machinery have been reported [17–20], plant cells have eukaryotic protein processing and post-translational modifications which are essential for glycoprotein production and proper protein folding. Moreover, inadvertent contamination with animal-derived pathogens can be avoided [21]. In addition, use of RAmy3D promoter and its signal peptide leads to the secretion of the target protein into rice cell culture media which aids downstream purification[22]

Natural mannose phosphorylation in plants has not been reported but recombinant α/β subunits of mammalian UDP-N-acetylglucosamine:glycoprotein N-acetylglucosamine-1-phosphotransferase (GNPTAB) and N-acetylglucosamine-1-phosphodiester alpha-N-acetylglucosaminidase (NAGPA) from rat liver, simple eukaryotes and humans can mediate this reaction *in vitro* [23–26]. N-glycans that have more than six mannose residues were found to be suitable substrates for this enzyme. In previous attempts to produce rhGAA from rice cell suspension culture, rhGAA was highly expressed in both wild-type and N-acetylglucosaminyltransferase-I (gntI) mutant rice cells and its α-glucosidase activity was highly similar to that of rhGAA from CHO cells [27, 28]. However, its N-glycan pattern, plant-specific or Man5 type, that are the most abundant in each line, were an unfavorable substrate for enzymatic *in vitro* phosphorylation [26].

The *α*-mannosidase I in ER and/or cis-Golgi apparatus conducts mannose trimming in the early stage of the N-glycosylation pathway [29]. For more detail, sequential removal of glucose residues (Glc) from initial form of N-glycan, Glc_3_Man_9_GlcNAc_2_Asn by *α*-glucosidases I and II yield Man_9_GlcNAc_2_. The *α*-mannosidase I in ER remove the terminal *α*1-2Man from the central arm of Man_9_GlcNAc_2_ and *α*-mannosidase I in cis-Golgi apparatus two Man residues of the Glc*α*1-3Man*α*1-2Man*α*1-2 moiety from some of N-glycans in the cis-Golgi retain a glucose residue because of incomplete processing in the ER, thereby both generating a Man8GlcNAc2 isomer by using different mechanism. In Arabidopsis, inhibition of ER-type *α*-mannosidase I or two Golgi-*α*-mannosidase I blocked N-glycan maturation and resulted in Man9 and Man8 rather than Man5, respectively [30]. Through phylogenetic analysis, Os04g0606400 in rice was expected to encode a putative Golgi-*α*-mannosidase I. In a more recent study, use of two different inhibitors of *α*-mannosidase I in the ER and/or cis-Golgi apparatus (kifunensine), and *α*-mannosidase II in the ER and/or medial-Golgi (Swainsonine) during the production of rhGAA in wild type rice successfully achieved production of rhGAA with Man7-9 type of N-glycan [31]. These studies show that genetic mutation of a-mannosidase I or pharmacological inhibition results in the production of rhGAA with Man7-9 type N-glycan.

In this study, I screened T-DNA insertional mutant rice expected to be defective in putative Golgi-*α*-mannosidase I genes (Δ*α-man*I). This Δ*α-man*I mutant rice showed highly decreased mannose trimming activity and was used for the production of rhGAA with mannosidic N-glycans.

## Materials and Methods

### Vector construction and Agrobacteria transformation

The rice expression vector pMYD85, carrying rhGAA gene, in which protein expression is induced by the rice *α*-amylase 3D promoter under the sugar starvation conditions, was constructed in previous study [27, 28]. For the purification, the signal sequence of rice *α*-amylase 3D (3Dsp), with 8 histidine or FLAG tags and enterokinase cleavage sequence were fused to N-terminus of pre-mature hGAA (57-952 a.a). In detail, 3Dsp was amplified by primer pair 3Dsp F1(XbaI): 5’-TTT CTA GAA TCA GTA GTG GTT AGC -3’ and GAA(8His-EK) R1: 5’-CTT GTC ATC ATC GTC GTG GTG ATG GTG ATG GTG ATG GTG GTG AGC TGG GTG AGT CTC CTC CA -3’ for the addition of a his tag, primer pair of 3Dsp F1(XbaI) and GAA(FLAG) R1: 5’-CTT GTC ATC ATC GTC CTT GTA GTC GTG AGC TGG GTG AGT CTC CTC CA-3’ for FLAG tagging. The partial hGAA sequence was amplified by primer pair EK-GAA F1: 5’-GAC GAT GAT GAC AAG CAG CAG GGA GCC AGC AGA CCA GGG CCC CG -3 ‘ and hGAA partial (StuI) R1: 5’-TTA GGC CTG TGA TAT ACT GCG AGG GCA GCG AGG -3’. These DNA fragments were fused by overlapping PCR using primer pair 3Dsp F1 (XbaI) and hGAA partial (StuI) R1 and then N-terminus of pMYD85 was replaced using XbaI+StuI restriction enzyme site to form pMYD532 and pMYD534. These vectors were transformed into Agrobacterium EHA-105 for rice transformation using the helper plasmid pRK2013 by tri-parental mating method [32].

### Screening of *α*-mannosidase I mutant candidates

T-DNA insertional mutant rice seeds of *α*-mannosidase I candidates for rhGAA expression were obtained from RiceGE (http://signal.salk.edu/cgi-bin/RiceGE). Calli were produced from 16 individual T-DNA mutant lines consisting of nine lines for chromosome 1, single lines for chromosome 2, two lines for chromosome 3, three lines for chromosome 4 and a single line for chromosome 5. Cell line showing reduced intensity or no bands in western blot using rabbit anti-horseradish peroxidase antibody (P7899, Sigma-Aldrich) which can recognize core *α*1,3-fucose and β1,2-xylose were propagated and used for further analysis (supplementary figure 1) [33–36].

Genotyping PCR to confirm T-DNA insertion into the gene of Os04g0606400 by using primer pairs LP (5’-TTGCAATGCCTACTGAGTGG-3 ‘) and LBP (5’-GTC GAG AAT TCA GTA CAT T-3’), or RP (5’-CTCTCGGGAGTTCTCACTGG-3’) and RBP (5’-TTG GGG TTT CTA CAG GAC GTA AC-3’) were conducted under the following PCR conditions: 1 cycle at 94 °C for 5 min; 30 cycles at 94 °C for 30 s, 55 °C for 30 s, and 72 °C for 45 s, followed by 1 cycle at 72 °C for 5 min. Homozygous T-DNA insertion was negatively identified by PCR using primer pair LP and RP under same condition (Figure 1A-B). Cell line shows no band which means homozygous T-DNA insertion was selected and used for further analysis.

**Figure 1.**
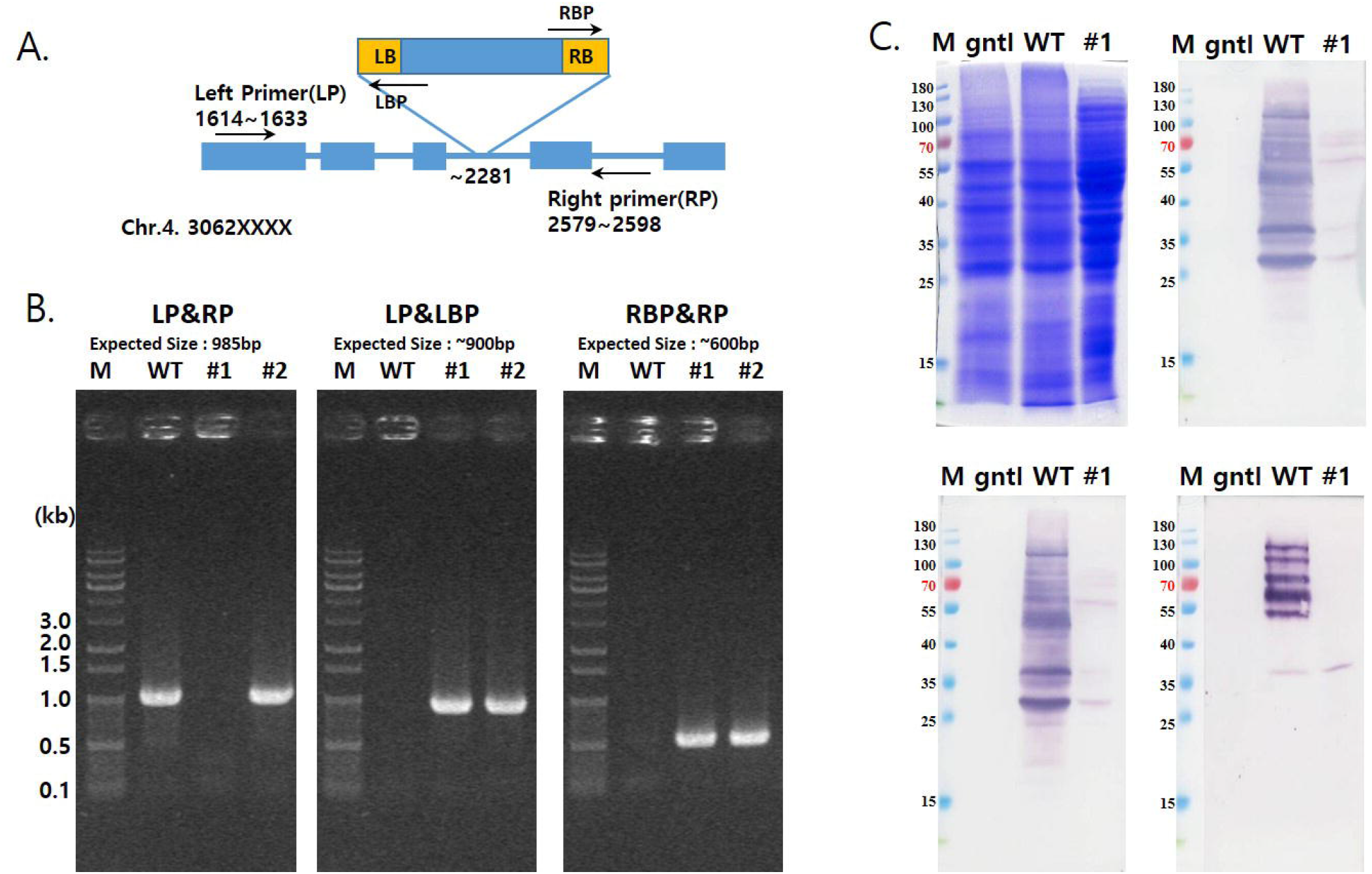
Characterization of Δ*α-man*I mutant rice. (A) Scheme of genotyping PCR. Depends on genome sequence of rice (Acession number. AP014960.1) and information from mutant rice library, genotyping PCR primers were designed. Numbers in the figure mean position on rice chromosome 4 (e.g 30621614-30621633 for Left primer) (B) Agarose gel images of genotyping PCR. Genomic DNA was extracted from putative Δ*α-man*I T-DNA insertional mutants and used for PCR. Lane M, 1kbp plus 100 DNA ladder (Elpis); Lane WT, wild type rice; Lane #1, putative homozygote; Lane #2, putative Heterozygote. (C) SDS-PAGE and Western blot analysis using antibodies against to plant specific N-glycans. Protein was extracted from calli and separated by SDS-PAGE(left upper), and then transblotted to nitrocellulose membrane to predict if plant specific N-glycans such as *α*1,3-Fucose(right upper), β1,2-xylose (left lower) and Lewis a epitopes (right lower). Lane M, Pre-stained protein ladder; Lane gntI, gntI mutant rice callus extracts; Lane WT, wild-type rice callus extracts; Lane 1, Callus extracts from #1 line in fig.1B.

Again, protein extracts from wild type, *gnt*I mutant (SAI2G12, http://signal.salk.edu/cgi-bin/RiceGE) [27] and homozygous T-DNA insertional mutant rice above were loaded to 10% (w/v) SDS-polyacrylamide gel and separated by electrophoresis. The protein bands were visualized by Coomassie brilliant blue. For western blot, proteins were transferred to Nitrocellulose membrane (Hybond-C, GE healthcare) in transfer buffer (50 mM Tris, 40 mM glycine, and 20% methanol) using a Trans-Blot^®^ SD Semi-Dry Transfer Cell (Bio-Rad) at 25 V for 30 min. Membranes were blocked with 10% (w/v) skim milk in Tris-Buffered Saline with Tween-20 on a rotary shaker overnight. *α*1,3-fucose and β1,2-xylose were detected by 1:5000 dilution of primary antibodies of *α*1,3-fucose (Agrisera), β1,2-xylose (Agrisera, Sweden), followed by 1/7000 dilution of anti-rabbit IgG alkaline phosphatase conjugate (Sigma-Aldrich) as secondary antibody. And Lewis a epitope was detected by 1:200 dilution of JIM84 antibody[37], followed by 1/7000 dilution of anti-rat IgG alkaline phosphatase conjugate (Sigma-Aldrich). Color was developed using BCIP/NBT (Millipore). Reaction was stopped within 1 min by washing the membrane with distilled water.

### Callus induction and rice transformation

Rice seeds were dehusked and washed several times with 70% (v/v) ethanol, 2% (v/v) NaOCl and autoclaved distilled water, respectively. The sterilized seeds were germinated for one week on callus induction agar plate (N6 plates) containing 4 g/L of CHU salts, 30 g/L of sucrose, 2 mg/L of 2,4-D, 0.2 mg/L of kinetin, and 4 g/L phytagel with 50mg/L hygromycin B as selection marker.

Generated calli were isolated, propagated and used for western blot or genotyping PCR for screening of Δ*α-man*I mutants. Putative Δ*α-man*I mutant rice calli were further propagated and used for *A.tumefaciens-mediated* transformation with pMYD85, pMYD532 and pMYD534, respectively. Infected calli were cultured on N6 plates without antibiotics for 3 days in dark condition and washed with distilled water containing 500 mg/L of Cefotaxime and transferred to selection media (N6 plates supplemented with DL-phosphinothricin (PPT; 4.4 ppm)) After 3-4 weeks, transgenic calli that survived were moved onto new selection media and propagated.

### Genomic DNA extraction and PCR

To confirm hGAA gene integration into the genome of Δ*α-man*I mutant rice, genomic DNA PCR was conducted after extraction of genomic DNA using a Zymobead genomic DNA kit (Zymo Research). Primer pair amplifying hGAA and His-hGAA (GAA gD F1: 5’-GTT CCC CGA GAG CTG AGT GG-3’ and GAA gD R1: 5’-GGT AGA AAG GGT GAG ACC CG -3’) was used for PCR reaction. Thermal cycling was performed for 30 cycles, consisting of 1 min at 94°C, 1 min at 55°C, and 3 min at 72°C. The PCR products were electrophoresed on a 1.0% (w/v) agarose gel, visualized by staining with ethidium bromide, and observed under UV light.

### Rice cell suspension culture and induction of protein expression

Δ*α-man*I rice mutant and Δ*α-man*I transformed with individual pMYD85 (D85), pMYD532 (D532) and pMYD534 (D534) were cultured in 50 mL N6 medium containing 2 mg/L 2,4-D, 0.02 mg/L kinetin, 3% (w/v) sucrose, and 50 mg/L hygromycin B in 300 mL flask at 28 °C in the dark using a rotary shaker with a rotation speed of 100 rpm. 10 mL of Δ*α-man*I rice mutant cells, transferred every 3-4 days and spent media were collected and kept in −70 °C for further analysis.

To induce target gene expression under control of the Ramy3D promoter, 5 g of fresh cells collected by aspiration, washed with distilled water and inoculated into 50 mL fresh N6 (CHU) medium without sucrose. At 9 days after sugar starvation, spent media was harvested and store in - 70 °C for further analysis. The rhGAA expression levels of each cell line were measured by comparing band intensity with that of positive control on SDS-PAGE gel using Image J software.

### Purification of rhGAA

rhGAA secreted from D532 was purified using 1 mL Ni-NTA agarose resin (Qiagen) according to the manufacturer’s instructions and as described previously [27] with slight modification. Briefly, 0.45 μm filtered 50 mL of culture media produced under sugar starvation conditions was loaded. The unbound protein was washed with 10 mL of 1X Phosphate-buffered Saline (PBS), pH 7.0, and the bound protein was eluted with 1 mL of PBS (pH 4.3) [27] and neutralized using 25 μL of 1 M Tris-HCl, pH 8.0.

rhGAA secreted from D534 was also purified using 1 mL of M2 anti-FLAG affinity column (GE healthcare) according to the manufacturer’s instruction. 30 mL of culture media under sugar starvation was loaded to prepared resin and flow-through was collected every 10 mL. The resin was washed with 5 column volumes of TBS (50 mM Tris HCl, with 150 mM NaCl, pH 7.4) to remove unbound protein, and followed by elution using 1 mL aliquots of 0.1 M glycine-HCl, pH 3.5, into vials containing 25 μL of 1 M Tris-HCl, pH 8.0.

### SDS-PAGE and western blot

To detect rhGAA in the samples, SDS-PAGE and western blotting were used. 30 μL of each samples were loaded to 10% (w/v) SDS-polyacrylamide gel and separated by electrophoresis. The protein bands were visualized by Coomassie brilliant blue. For western blot, proteins were transferred to Nitrocellulose membrane (Hybond-C, GE healthcare) in transfer buffer using a Trans-Blot^®^ SD Semi-Dry Transfer Cell (Bio-Rad) at 25 V for 30 min. Membranes were blocked with 10% (w/v) skim milk in Tris-Buffered Saline with Tween-20 on a rotary shaker overnight. Target protein was detected by 1/2000 dilution of rat anti-GAA polyserum [27], followed by 1/7000 dilution of anti-rat IgG alkaline phosphatase conjugate (Sigma-Aldrich) as secondary antibody. Colour was developed using BCIP/NBT (Millipore). Reaction was stopped within 5 min by washing the membrane with distilled water.

### MALDI-TOF

The collected media from Δ*α-man*I was centrifuged at 15,000g for 30 min at 4 C. The supernatant was lyophilized to reduce the volume and dialyzed in the distilled water. After filtration by 0.45 μm filter, it was applied to a Con A-Sepharose 4 B column (GE Healthcare), and glycoproteins were eluted with 0.5 M *α*-D-methylglucoside. This sample and purified Δ*a-man*I-GAA (D532) prepared above were lyophilized, each sample (2 mg) was digested with 50 μg trypsin (T0303; Sigma-Aldrich) and chymotrypsin (C4129; Sigma-Aldrich) at 37 °C for 16 h and boiled for 10 min for inactivation of enzyme. N-glycans were released by incubation with 1.0 mU PNGase A (Roche Diagnostics) in 60 μL 0.5 M citrate/phosphate buffer at pH 4.0 for 16 h and purified using a Carbograph Ultra Clean column (Alltech) as described previously [38]. Pyridylamination of N-glycans was performed using a pyridylamination manual kit (Takara Bio Inc.) according to the manufacturer’s instructions. For further purification of the 2-aminopyridine (PA)-N-glycans, NP-high performance liquid chromatography (HPLC) was performed using an TSK gel Amide-80 column (4.6× 250 mm; Tosoh Co.) and a Waters 2690 Alliance HPLC separation module (Waters Co.) equipped with the 474 Fluorescence Detector (Waters Co.), as described previously [39]. Fluorometric detection was carried out at excitation and emission wavelengths of 310 and 380 nm, respectively. The molecular masses of PA-N-glycans were determined using the 4800 Plus matrix-assisted laser desorption ionization time-of-flight/time-of-flight (MALDI-TOF/TOF) analyzer (Applied Biosystems). The PA-glycans purified from the pooled NP-HPLC fractions (75–130 min) were subjected to MALDI-TOF/MS analysis as described previously [40]. The mass spectra were collected in the reflector positive ion mode using 2,5-dihydroxybenzoic acid (10 mg/mL in 50% acetonitrile/0.1% trifluoroacetic acid and 10 mM NaCl) as a matrix. Purified PA-N-glycans were mixed with an equal volume of matrix solution, and 2 μL of the mixture were placed on the target plate and air-dried [41]. An m/z range of 1000–2400 was measured and analysed using DATA Explore software (Applied Biosystems). Peaks were assigned according to the calculated molecular masses [M+Na+] of PA-N-glycans, and mannose, galactose, fucose, xylose, and N-acetylglucosamine residues were identified. N-glycan structures were assigned by comparison with the plant N-glycans identified to date using SimGlycan version 2.8 (PREMIER Biosoft Int.). Relative abundances were calculated based on the areas of the corresponding peaks in the mass spectra, as described previously [42].

## Results

### Identification of Δ*α-man*I mutant

To select the appropriate Δ*α-man*I mutant rice line for expression of rhGAA with high mannose N-glycans, α-mannosidase I in rice known to be localized in the ER or Golgi and containing the glycosyl hydrolase family 47 domain which catalyse mannose trimming reaction were searched from protein database (https://www.uniprot.org/) and rice seeds have T-DNA insertion on those position were purchased through RiceGE: Rice Functional Genomic Express Database (http://signal.salk.edu/cgi-bin/RiceGE) from Kyung Hee University, Republic of Korea and Huazhong Agricultural University, China (supplementary table 1.)[30]. Calli were induced from the mutant rice seeds have T-DNA insertions on five different putative *α*-mannosidase I gene in rice genome, respectively. Protein extracts from these putative rice calli lines were analysed by western blot using anti-HRP antibody for screening of mutant lacking *α*1,3-fucose and β1,2-xylose which reflect the blockage of N-glycan maturation. Rice mutant possessing T-DNA insertions in Os04g0606400 (LOC_Os04g51690, PFG-3A-12868) showed highly reduced bands correspond to *α*1,3-fucose and β1,2-xylose was used for further analysis (supplementary figure 1.). Homozygous and heterozygous T-DNA insertional pattern were observed from two individual Δ*α-man*I mutants by genotyping PCR using LP, RP, LBP and RBP (Fig.1A and 1B). #1 and #2 those are individual rice mutant of Os04g0606400 showed band from PCR using primer pair LP and LBP, or RP and RBP which means T-DNA insertion on expected position of rice genome. Homozygous T-DNA insertion was confirmed by PCR using primer pair LP and RP targeting rice genome flaking T-DNA insertion site. Homozygote (#1) shows no band because of its large length of T-DNA (~8 kb) while wild type and heterozygote (#2) show amplicon. Amplified DNA fragments from #1 were extracted from Agarose gel and analysed by DNA sequencing (data not shown), revealing T-DNA insertion into the intron between the 11^th^ and 12^th^ exon as expected. The growth rate of this homozygous Δ*α-man*I mutant rice callus was similar to that of the wild type without any severe defects even under sugar starvation. For further analysis, western blot using anti-*α*1,3-fucose, anti-β1,2-xylose and JIM84 antibody showed highly reduced band intensity, respectively (Fig. 1C).

This demonstrates that the Δ*α-man*I mutation inhibits N-glycan maturation such as attachment of plant N-glycan species, *α*1,3-fucose, β1,2-xylose, β1,3-galactose and *α*1,4-fucose in rice.

### Expression and purification of Δ*α-man*I-GAA

Calli were transformed with Agrobacteria containing the vectors pMYD85 (rhGAA without tag), pMYD532 (8His+EK+rhGAA) or pMYD534 (FLAG-EK-rhGAA) were screened by genomic DNA PCR. Eight or more cell lines from each construct were chosen for propagation and induced to express rhGAA under sugar starvation conditions. D85#7, D532#3 and D534#16 transformed with individual pMYD85, pMYD532 and pMYD534 showed the highest expression of premature form Δ*α-man*I-GAA (~110 kDa) at the 9 days after sugar starvation on SDS-PAGE and Western blot, respectively (Fig. 2). The rhGAA expression levels of each pMYD85#7, pMYD532#3 and pMYD534#16 cell line were approximately 10 mg/L, 21mg/L and 12 mg/L by comparing band intensity with positive control on SDS-PAGE.

**Figure 2.**
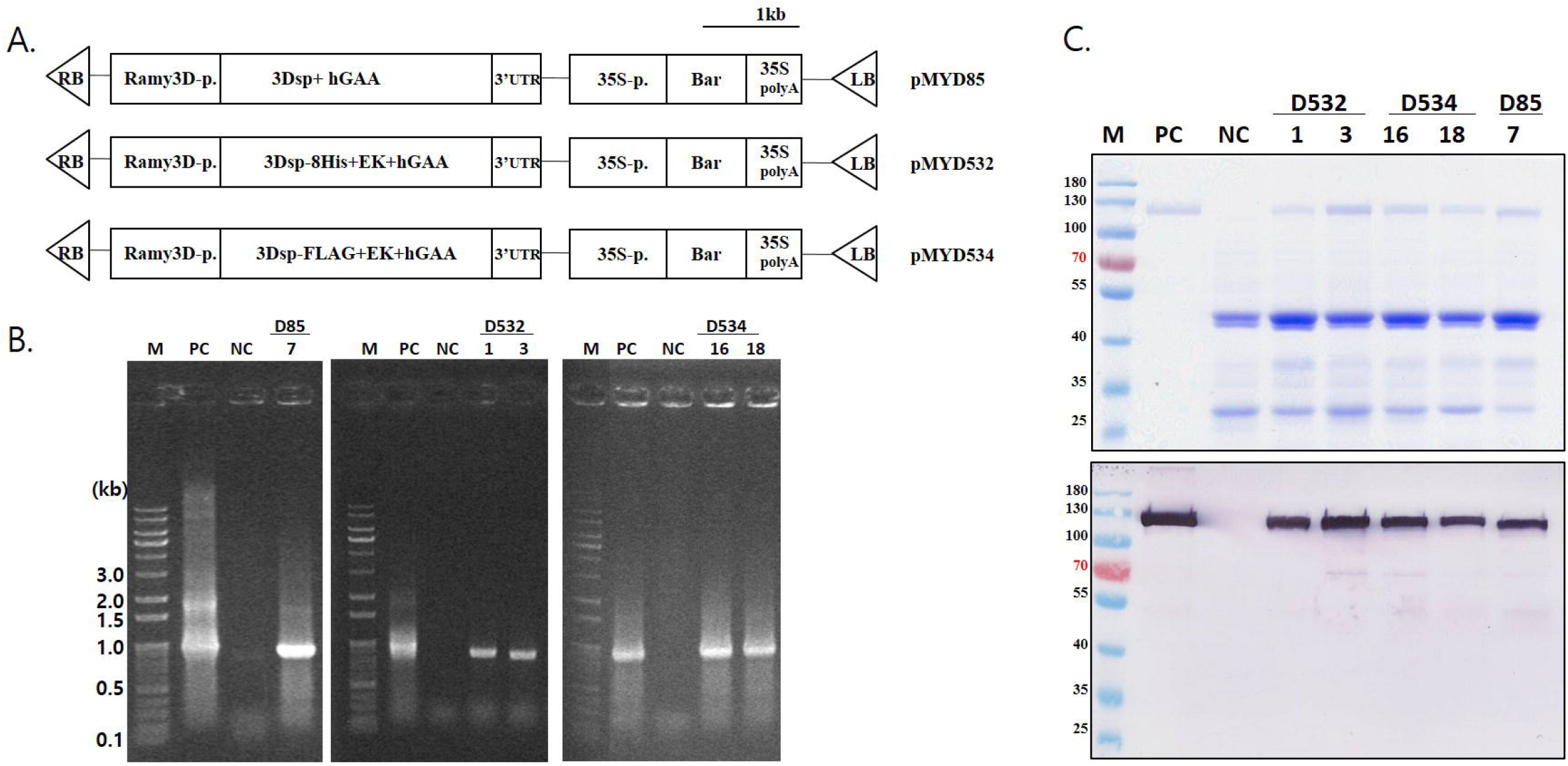
Expression of rhGAA in Δ*α-man*I mutant rice. (A) Rice expression vectors construction. Rice expression vectors for the human acid *α*-glucosidase (hGAA) gene (pMYD85), hGAA with six histidine residues (pMYD532), or hGAA with FLAG-tag on N-terminus with the signal peptide of the rice *α*-amylase 3D gene and its own pro-peptide, located between the rice *α*-amylase 3D promoter (Ramy3D-p) and the 3’ untranslated region (3’ UTR). LB, T-DNA right border; 35S-p, CaMV 35S promoter; Bar, bialaphos resistance gene; 35S polyA, terminator of 35S gene; RB, T-DNA right border; EK, Enterokinase cleavage sequence between purification tags and hGAA. (B) Agarose gel images of genomic DNA PCR. Lane M, 1kbp plus 100 DNA ladder (Elpis); PC, Plasmid DNA; NC, genomic DNA PCR product of wild type rice; Lane numbered, genomic DNA PCR products of putative transgenic calli. (C) Rice cell suspension culture media 9 days after induction (dai). SDS-PAGE (upper) and Western blot using rat anti-GAA antiserum (lower). Lane M, Pre-stained protein ladder; Lane PC, CHO cell derived rhGAA 500ng; Lane NC, wildtype rice culture media under sugar starvation; Lane numbered, putative transgenic rice culture media under sugar starvation.

In the case of D532 and D534, expressed rhGAA was purified by affinity chromatography using the eight histidine residues or FLAG tag on N-terminus (Fig. 3). Histidine tagged Δ*α-man*I-GAA could be purified successfully with protocol used for purification of rhGAA with histidine tag on C-terminus in previous studies [27, 28]. N-terminal FLAG tagged Δ*α-man*I-GAA also could be highly purified using anti-FLAG resin but showed lower binding capacity. Impurities on SDS-PAGE and Western blot might be matured form of rhGAA by acidic condition during purification or host cell protein unspecifically bound. Hence, Histidine tagged Δ*α-man*I-GAA was used for further analysis.

**Figure 3.**
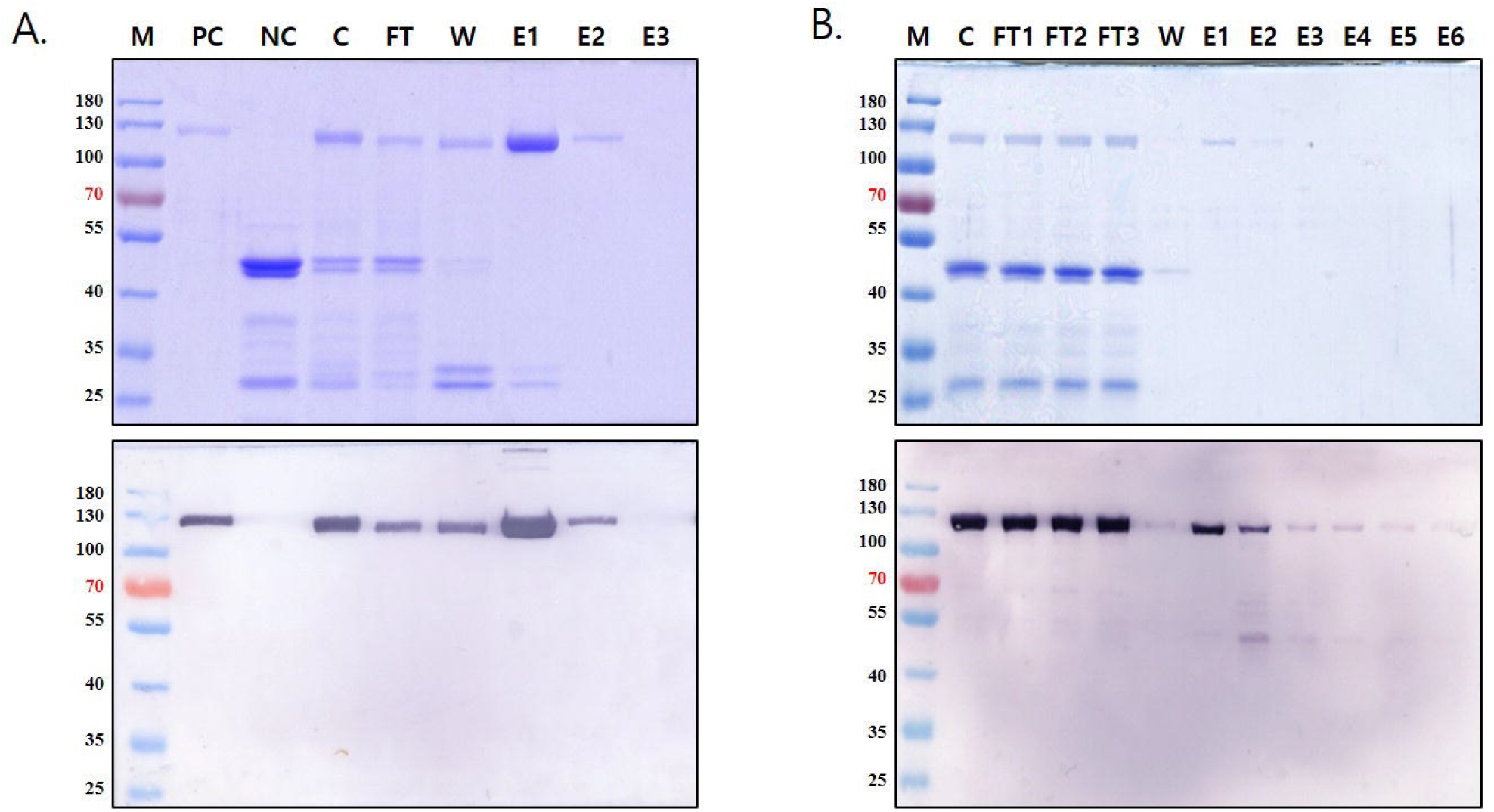
Purification of recombinant human acid *α*-glucosidase (rhGAA) from rice cell suspension culture. SDS-PAGE and Western blot using anti-GAA antibody of purified rhGAA by affinity chromatography using N-terminus histidine tag (A) or FLAG tag (B). Lane M, Pre-stained protein ladder; Lane PC, CHO cell derived rhGAA 500ng; Lane NC, Δ*α-man*I mutant rice cultured media under sugar starvation; Lane C, crude cultured media under sugar starvation; FT, Flow through from the affinity column; Lane W, Wash from the affinity column; Lane E, eluted fractions.

### N-glycan profiling

N-glycan patterns of glycoproteins in culture media of Δ*α-man*I mutant and purified Δ*α-man*I-GAA were analysed and compared to that of wild-type rice (Fig.4 and table. 1). N-glycans from Δ*α-man*I mutant and Δ*α-man*I-GAA were almost all high mannose type of N-glycans while wild type cell lines shows typical pattern of plant N-glycan contents. Interestingly, the most abundant N-glycan from Δ*α-man*I was Man7 (48.9%), followed by Man8 (33.3%), Man6 (7.0%) while that of Δ*α-man*I-GAA was Man8 (63.8%), Man7 (22.5%) and Man6 (5.4%). This difference may be because various host glycoproteins were present in the culture media of the Δ*α-man*I mutant while Δ*a-man*I-GAA was specifically purified. Most importantly, N-glycan patterns in both Δ*α-man*I and Δ*α-man*I-GAA reflect that gene disruption of Os04g0606400 highly inhibited N-glycan maturation in rice.

**Figure 4.**
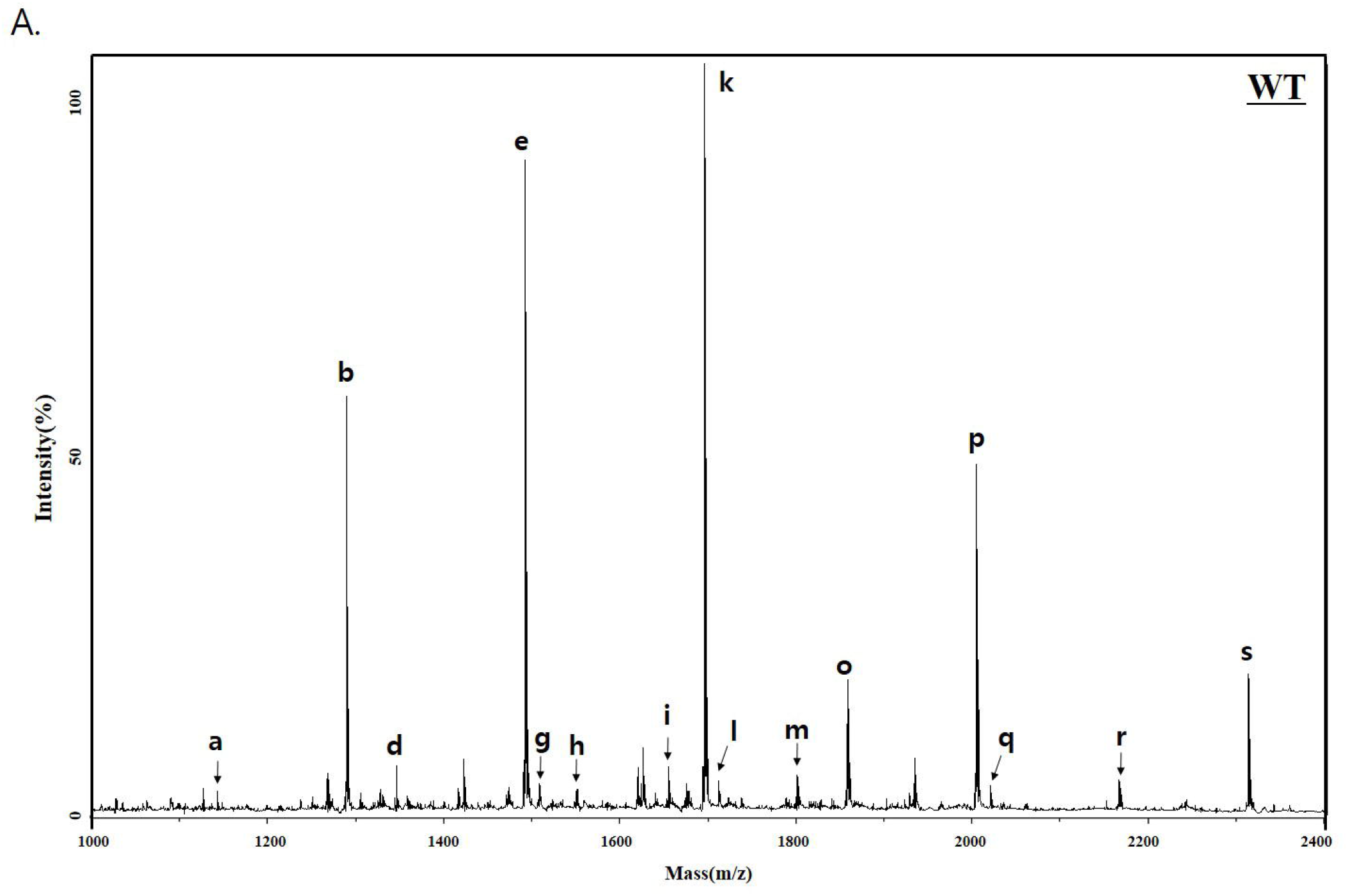
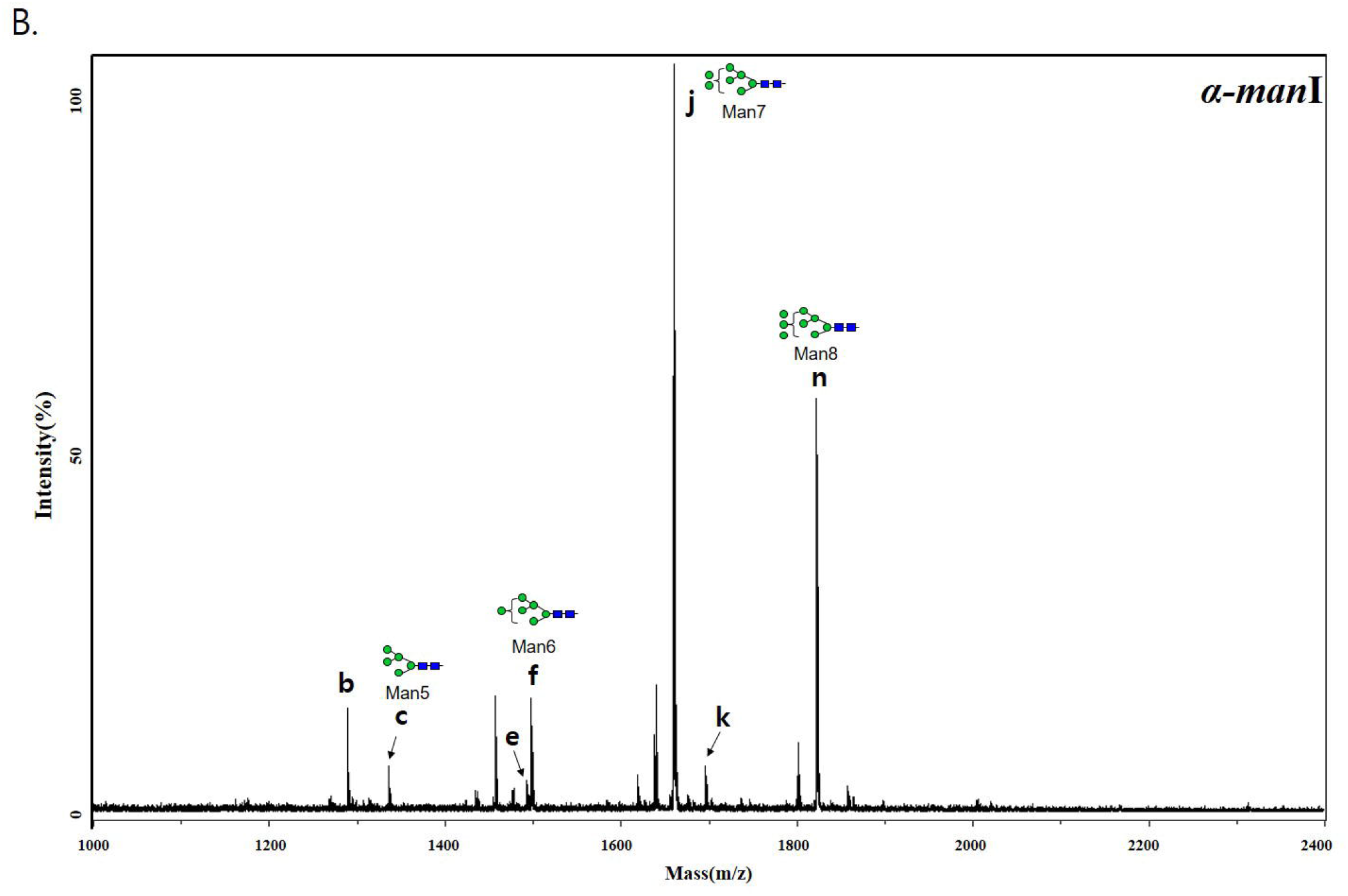
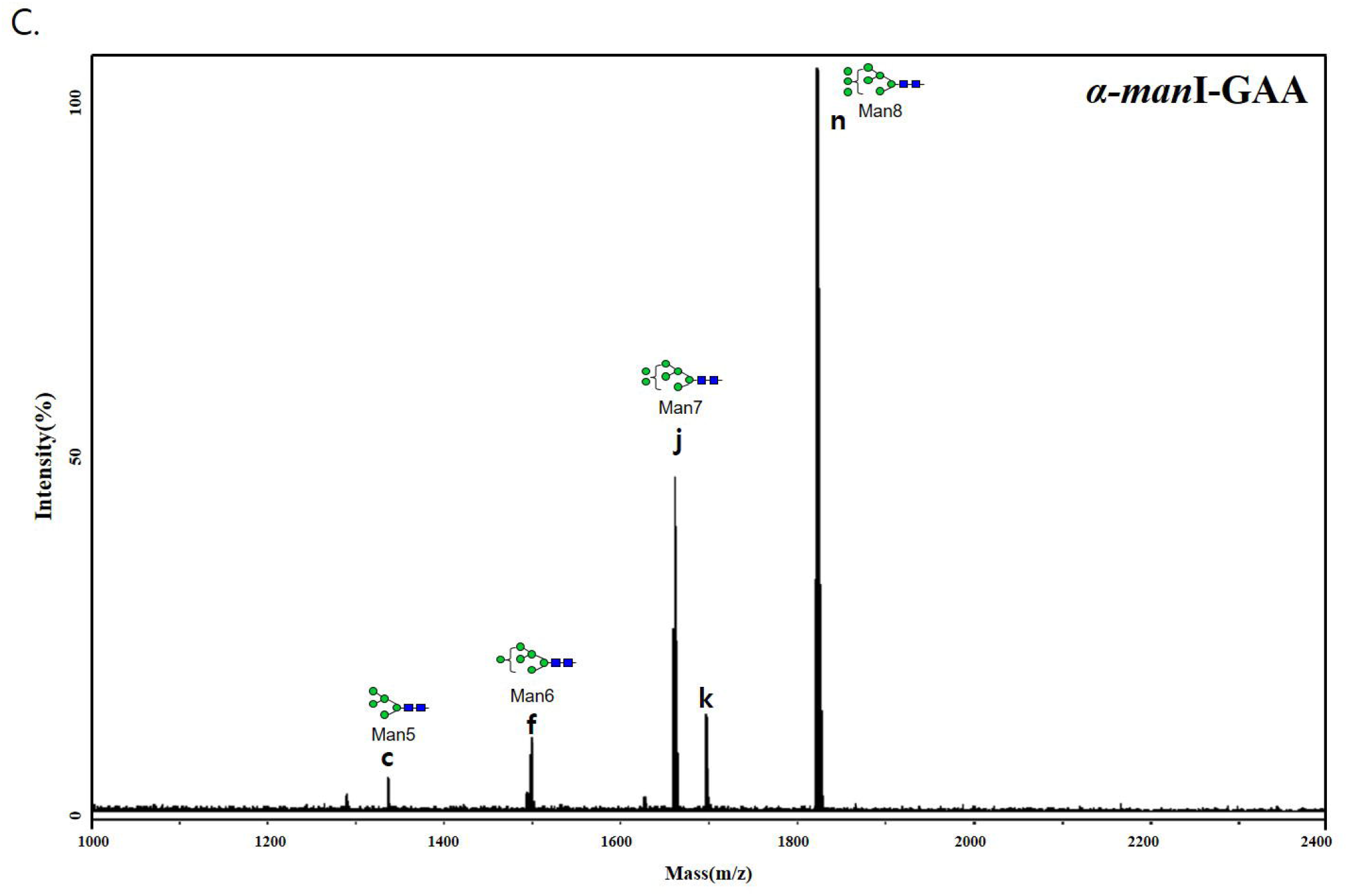
Matrix-assisted laser desorption ionization time-of-flight/time-of-flight (MALDITOF)/mass spectra of 2-aminopyridine (PA)-N-glycans released from extra cellular glycoproteins. Peaks from each sample were characterized in the positive reflector ion mode, and detected ions were assigned to Na-adducts (M+Na+) of the PA-N-glycans. Glycoproteins were collected from the culture media from wild-type rice (A), Δ*α-man*I mutant rice (B), and purified recombinant Δ*α-man*I-GAA (C), respectively.

**Table 1.**
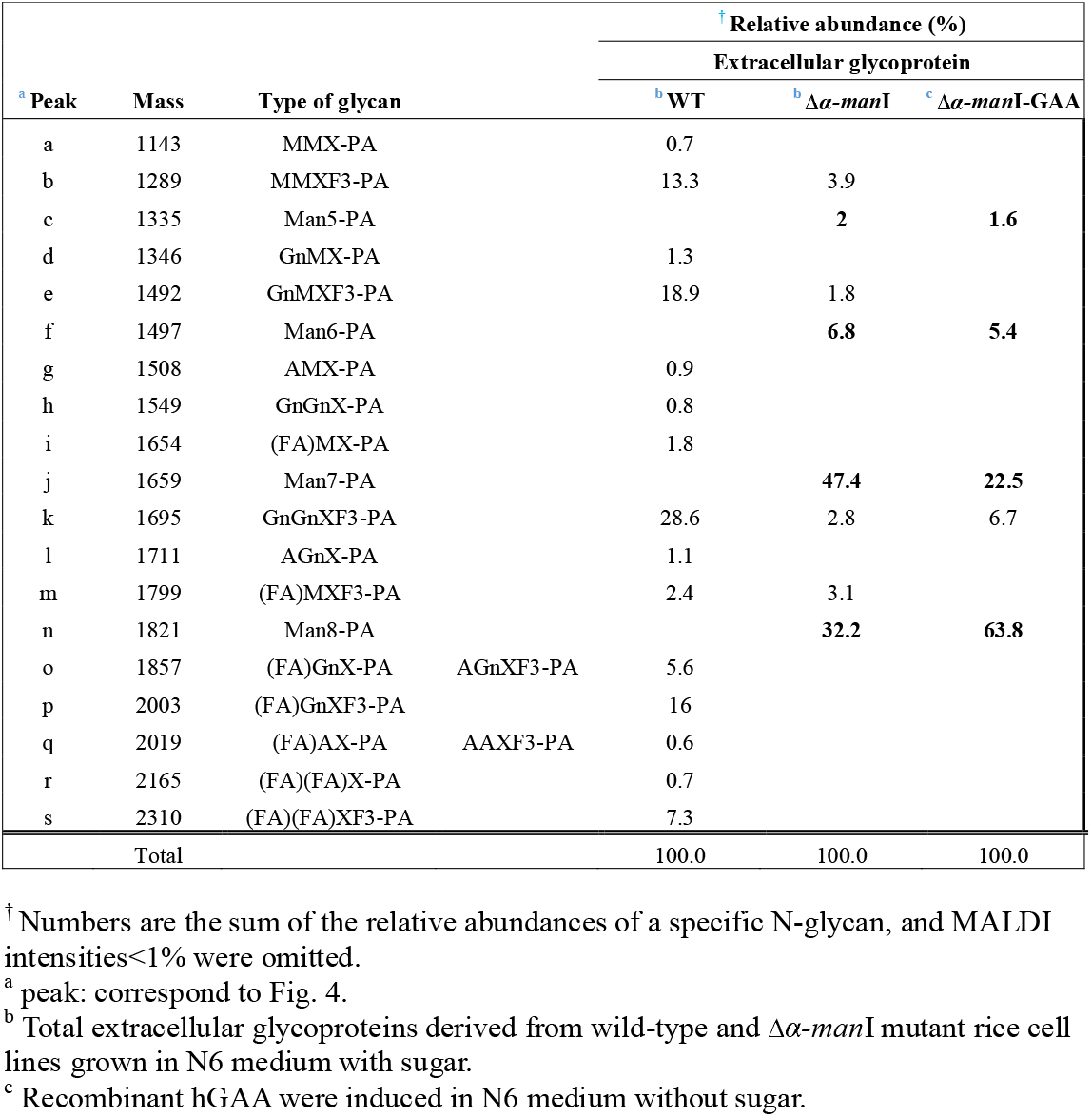
Comparison of N-glycan patterns among total secreted glycoproteins from wild type and Δ*α-man*I mutant rice and purified Δ*α-man*I-GAA.

## Discussion

In this study, I suggest a major role for Os04g0606400 in the early stage of plant N-glycan maturation and the successful expression of Δ*α-man*I-GAA. The main N-glycans on this Δ*α-man*I-GAA was Man8 which is known as suitable substrate for *in vitro* phosphorylation. Because the production of Δ*α-man*I-GAA in our study does not requires any inhibitor treatment, it can greatly reduce production costs and the cost burden for Pompe patients [31]. Although I used rhGAA in this study, other lysosomal enzyme that require M6P such as *α*-L-iduronidase[24], *α*-galactosidase A [43] can be produced using same strategy.

In screening of Δ*α-man*I mutant candidates, only the cell lines that have T-DNA insertions in Os04g0606400 showed highly decreased amounts of plant-specific N-glycans on western blot. However, it does not suggest that Os04g0606400 encodes the only enzyme that conducts N-glycan mannose trimming out of six candidates. Although other cell extracts showed bands corresponding plant specific N-glycans, genotyping PCR of all individual candidates was not conducted in this study. Furthermore, the seeds that I obtained from the library could be a mix of wild type, hetero- and homozygous mutant lines. In this study, I focused on screening for cell lines suitable for the production of rhGAA carrying more than six mannose residues on its N-glycan. Hence, I analysed only the Os04g0606400 mutant in detail and used it as the rhGAA expression host.

Because gntI mutant rice was unable to grow or regenerate in previous studies [27, 44], and Δ*α-man*I is located upstream of gntI in N-glycosylation pathway, I hypothesized possible lethality of the Δ*α-man*I knock-out mutant. Despite more than three attempts at plant regeneration from calli, no regenerated plant of the Os04g0606400 mutant were obtained while wild type callus were regenerated successfully (data not shown). I assume this mutation may causes a problem with plant regeneration and cell differentiation [44].

Both Δ*α-man*I-GAA tagged with Histidine and FLAG on N-terminus were successfully expressed and purified from transgenic Δ*α-man*I mutant rice. Although FLAG tag showed low binding to the affinity column, this can be improved by increasing volume of column. Furthermore, these tags would be be cleaved by enterokinase treatment while or after purification. By adding an enterokinase cleavage site between N-terminus tag and target protein, the heterogeneity of N-terminus, which can be made different signal peptide cleavage in plants and animals, would probably be improved.

Enzymatic activity was not measured in this study but rhGAA with 7-9 mannose residues produced from rice cell suspension culture under presence of mannosidase inhibitor showed similar *in vitro* enzymatic activity to that of CHO cell derived rhGAA in previous study [31]. Although this strategy blocks *α*-mannosidase I function at the protein level while Δ*α-man*I acts at the DNA level, both approaches target *α*-mannosidase I so it seems likely that Δ*α-man*I-GAA would probably have similar enzyme activity.

In N-glycan profiling, suitable N-glycan substrates for enzymatic *in vitro* phosphorylation, Man8, Man7, andMan6 were detected from rhGAA produced from Δ*α-man*I mutant rice. Interestingly, although N-glycan maturation is highly inhibited in this cell line, mannose trimming is not completely terminated with Man8 and processed further. Moreover, plant specific N-glycan species, GnGnXF3 (6.7% from Δ*α-manI-GAA,* 2.9% from Δ*α-man*I). GnMXF3 (1.9% from Δ*α-man*I) and MMXF3 (4.0% from Δ*α-man*I) were still detected. The reason of the presence of these unexpected N-glycans may be a result of insufficient knock-out by T-DNA insertion on intron or minor activity of *α*-mannosidase I isotype. This could be further characterized by using knock-out using other technique such as CRISPR/Cas9 in further studies. Nevertheless, it is clear that Os04g0606400 plays a major role in mannose trimming in N-glycosylation pathway in rice, and rhGAA expressed from this mutant mainly carried Man6-8 that are known as the most favorable substrates of *in vitro* phosphorylation using two enzyme, GNPTAB and NAGPA [45]. Although *in vitro* phosphorylation of Man6-8 on Δ*α-man*I-GAA was not tested in this study, studies involving *in vitro* phosphorylation of Man6-9 on the other recombinant lysosomal enzymes for ERT and the presence of Man6-8 on Δ*α-man*I-GAA suggest that Δ*α-man*I-GAA would also probably be phosphorylated via similar approaches [23, 24].

Therefore, this study provides knowledge about the early stage of N-glycosylation in plants and the production of rhGAA in Δ*α-man*I mutant rice. I expect that Δ*α-man*I mutant rice can be used for attractive expression host for production of lysosomal enzymes for ERT with cost effectiveness and safety.

## Acknowledgements

I acknowledge George P. Lomonossoff for English editing and supporting.

This research was supported by the Agriculture, Food and Rural Affairs Research Center Support Program of Ministry of Agriculture, Food and Rural Affairs (714001-7); Basic Science Research Program through the National Research Foundation of Korea (NRF) funded by the Ministry of Education (NRF-2018R1A6A3A01011033, NRF-2021R1A6A3A03038622); John Innes Centre, Knowledge Exchange and Commercialisation Innovation Funding (01IF 2020 GL01).

## Notes

### Competing Interest Statement

The authors have declared no competing interest.

