## supplemental figure.1 for "Production of recombinant human acid α-glucosidase with mannosidic N-glycans in α-mannosidase I mutant rice cell suspension culture"

### Slide 1
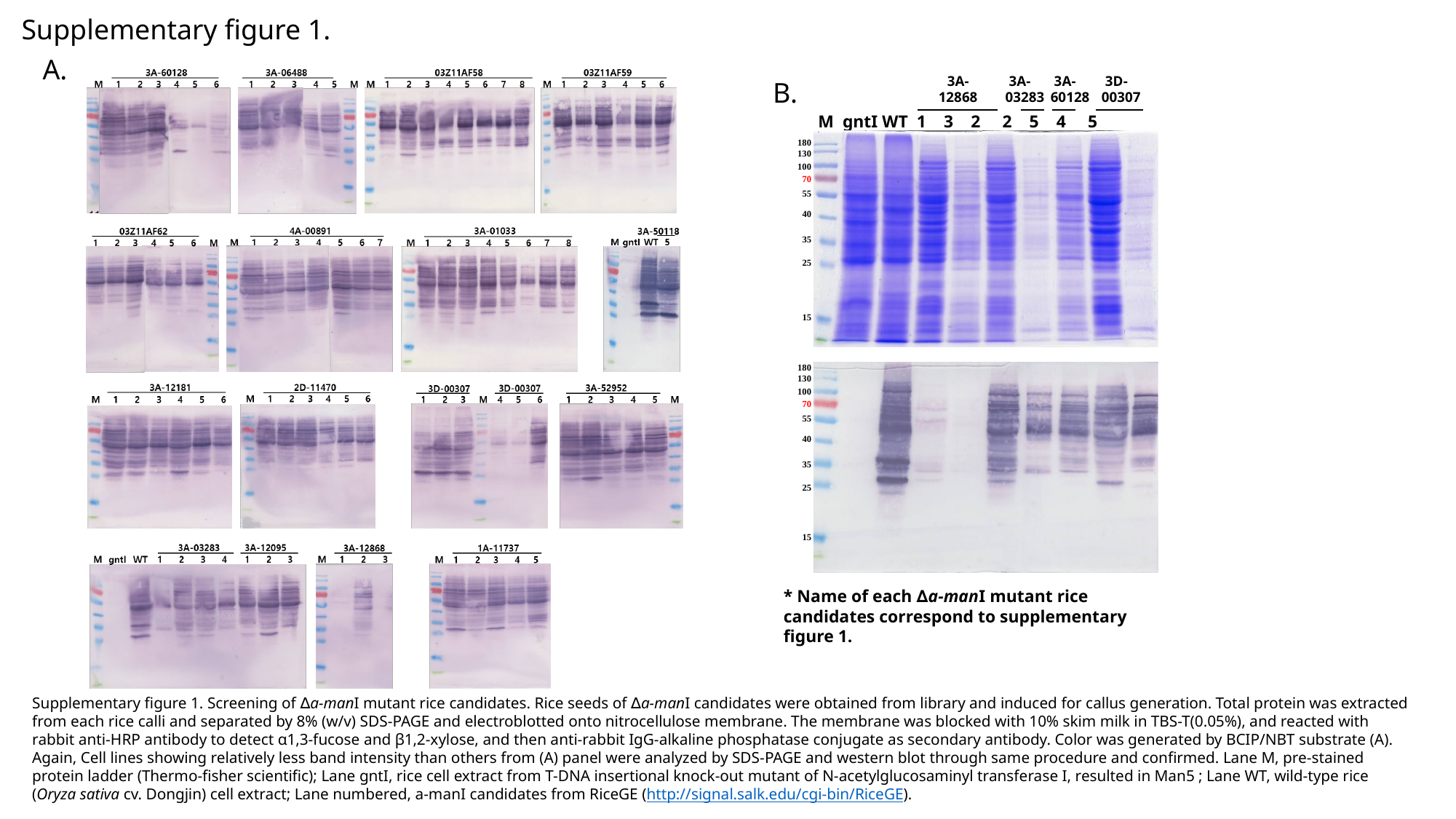

Supplementary figure 1.
A.
3A-
12868
 3A-
03283
 3A-60128
 3D-00307
B.
M gntI WT 1 3 2 2 5 4 5
180
130
100
70
55
40
35
25
15
180
130
100
70
55
40
35
25
15
* Name of each ∆a-manI mutant rice candidates correspond to supplementary figure 1.
Supplementary figure 1. Screening of ∆a-manI mutant rice candidates. Rice seeds of ∆a-manI candidates were obtained from library and induced for callus generation. Total protein was extracted from each rice calli and separated by 8% (w/v) SDS-PAGE and electroblotted onto nitrocellulose membrane. The membrane was blocked with 10% skim milk in TBS-T(0.05%), and reacted with rabbit anti-HRP antibody to detect α1,3-fucose and β1,2-xylose, and then anti-rabbit IgG-alkaline phosphatase conjugate as secondary antibody. Color was generated by BCIP/NBT substrate (A). Again, Cell lines showing relatively less band intensity than others from (A) panel were analyzed by SDS-PAGE and western blot through same procedure and confirmed. Lane M, pre-stained protein ladder (Thermo-fisher scientific); Lane gntI, rice cell extract from T-DNA insertional knock-out mutant of N-acetylglucosaminyl transferase I, resulted in Man5 ; Lane WT, wild-type rice (Oryza sativa cv. Dongjin) cell extract; Lane numbered, a-manI candidates from RiceGE (http://signal.salk.edu/cgi-bin/RiceGE).

### Slide 2
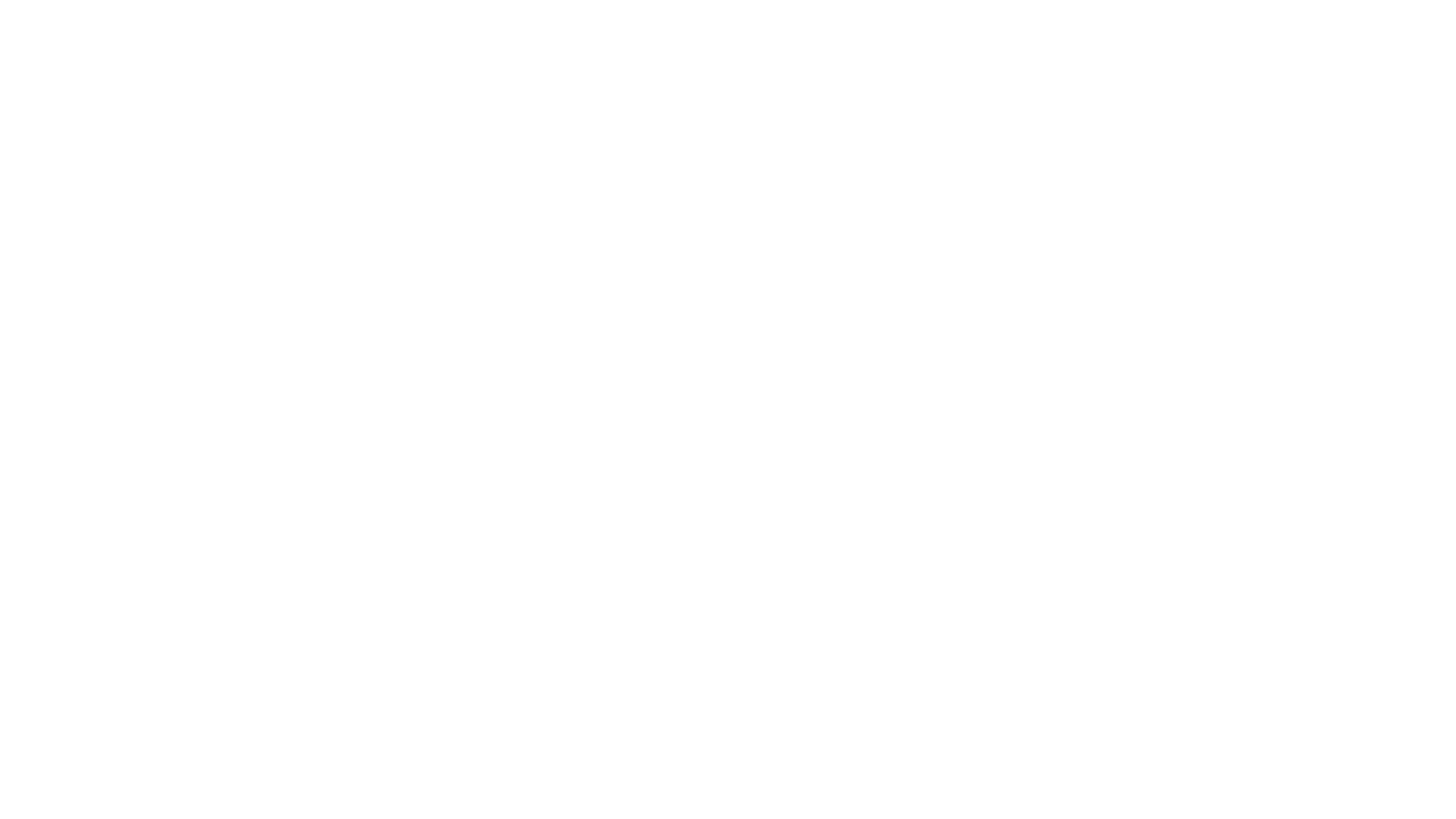
