## supplemental table.1 for "Production of recombinant human acid α-glucosidase with mannosidic N-glycans in α-mannosidase I mutant rice cell suspension culture"

**Supplementary table 1. ∆*α-man*I candidates screened for rhGAA expression in this study**

| **Gene locus** | **Line No.** | **Expected T-DNA Insertion site** | **Source** |
| --- | --- | --- | --- |
| LOC_Os01g56670 | PFG_3A-60128 | Intron | Kyung Hee University  (Republic of Korea) |
|  | PFG_3A-06488 | Intron |  |
|  | RMD_03Z11AF62 | Exon | Huazhong Agricultural University  (China) |
|  | RMD_03Z11AF58 | Exon |  |
|  | RMD_03Z11AF59 | Exon |  |
|  | PFG_3A-01033 | 300-UTR3 | Kyung Hee University  (Republic of Korea) |
|  | PFG_4A-00891 | Intron |  |
| LOC_Os01g13050 | PFG_3A-12181 | 1000-Promotor |  |
|  | PFG_2D-11470 | Exon |  |
| LOC_Os02g50780 | PFG_3A-50118 | Exon |  |
| LOC_Os03g19070 | PFG_3D-00307 | 1000-Promotor |  |
|  | PFG_3A-52952 | 300-UTR3 |  |
| LOC_Os04g51690 | PFG_3A-12868 | Intron |  |
|  | PFG_3A-12095 | 1000-Promotor |  |
|  | PFG_3A-03283 | 1000-Promotor |  |
| LOC_Os05g35266 | PFG_1A-11737 | 300-UTR5 |  |

Line numbers, gene locus and expected T-DNA insertion site were obtained from http://signal.salk.edu/cgi-bin/RiceGE
